# Elementary growth modes/vectors and minimal autocatalytic sets for kinetic/constraint-based models of cellular growth

**DOI:** 10.1101/2021.02.24.432769

**Authors:** Stefan Müller

**Affiliations:** Faculty of Mathematics, University of Vienna, Oskar-Morgenstern-Platz 1, 1090 Wien, Austria

**Keywords:** cellular self-fabrication, minimal pathways, growth rate, polyhedral geometry, elementary vectors, conformal generators

## Abstract

Elementary vectors are fundamental objects in polyhedral geometry. In metabolic pathway analysis, elementary vectors range from elementary flux modes (of the flux cone) and elementary flux vectors (of a flux polyhedron) via elementary conversion modes (of the conversion cone) to minimal cut sets (of a dual polyhedron) in computational strain design.

To better understand cellular phenotypes with optimal (or suboptimal) growth rate, we introduce and analyze classes of elementary vectors for models of cellular growth. *Growth modes* (GMs) only depend on stoichiometry, but not on growth rate or concentrations; they are elements of the growth cone. *Elementary* growth modes (EGMs) are conformally non-decomposable GMs; unlike elementary flux modes, they are not support-minimal, in general. Most importantly, every GM can be written as a conformal sum of EGMs. *Growth vectors* (GVs) and *elementary* growth vectors (EGVs) also depend on growth rate, concentrations, and linear constraints; they are elements of a growth polyhedron. Again, every GV can be written as a conformal sum of EGVs. To relate the new concepts to other branches of theory, we define *autocatalytic* GMs and the corresponding *(minimal) autocatalytic sets* of reactions.

As a case study, we consider whole cell models (simple kinetic models of self-fabrication). First, we use EGMs to derive an upper bound for growth rate that only depends on enzyme kinetics. Next, we study growth rate maximization (via control parameters for ribosome kinetics). In particular, we analyze *growth states* (GSs) and *elementary* growth states (EGSs) as introduced in [de Groot et al, 2020]. Unlike EGMs, EGSs depend on (metabolite) concentrations and growth rate. Most importantly, (i) we show that EGSs are support-minimal, (ii) we give a simple proof for the fact that maximum growth rate is attained at an EGS, and (iii) we show that, at every optimal EGS, the ribosome capacity constraint is active. Finally, we determine the dependence of EGSs on growth rate, and we study the relation between EGSs and minimal autocatalytic sets, EGMs, and elementary flux modes. Along the way, we point out (and resolve) mathematical issues in [de Groot et al, 2020].

## 1 Introduction

Cell proliferation involves cell growth and cell division. In more abstract terms, cellular *self-fabrication* involves cellular *self-maintenance*, including processes such as signaling, transport, metabolism, and gene regulation, and cellular *self-replication*, including macromolecular synthesis and cell cycle. Indeed, during one cycle, a cell self-fabricates all its constituents (metabolites, enzymes, lipids, DNA,…); it grows. By cell division, this leads to growth on the population level (microbial growth) or the tissue level (multicellular development).

The processes contributing to cellular growth can be summarized in a stoichiometric matrix *N* with rows corresponding to constituents (molecular species) and columns corresponding to processes (chemical reactions). In the deterministic setting, one has the dynamical system 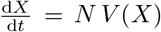 for the copy numbers *X* of molecules (extensive variables), as determined by *N* and the reaction rates *V* (*X*). After introducing concentrations *x* (intensive variables), one obtains 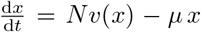, cf. [12]. The second term represents “dilution by growth”, where growth rate *μ* is given (as a function of *x*) by a linear constraint arising from dry weight or cell volume. The dynamical systems apply to individual cells or, as averages, to cell populations. Steady state for the intensive variables, that is, *x*(*t*) = *const* or *Nv*(*x*) = *μ x*, corresponds to balanced growth for the extensive variables, that is, *X*(*t*) = *X*(0) e^*μt*^. On the population level, one has 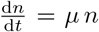 for the copy number *n* of cells, cf. [9, 22].

Particular models of cellular growth as well as model classes can be categorized by several dimensions: (i) the set of cellular processes modeled explicitly, (ii) the model structure – from detailed (elementary reaction steps as one extreme) to coarse-grained (one reaction for every process as another extreme), (iii) dynamic vs. steady-state, and (iv) kinetic (with nonlinear dependence *v*(*x*) of reaction rates on concentrations) vs. constraint-based (with linear constraints in *x* and *v*). For a recent review on mathematical models of cellular growth, see e.g. [6].

Traditional growth models often involve an approximative “biomass reaction”, which specifies macromolecular synthesis in terms of precursors. In terms of categories (i) and (ii), such models consider metabolism in detail and macromolecular synthesis as a coarse-grained process (the biomass reaction). In terms of (iii) and (iv), they often assume steady state and are constraint-based. In detail, concentrations *x* are fixed, and steady-state reaction rates (fluxes) *v* are considered as independent variables. The stoichiometric matrix has rows corresponding to metabolites and columns corresponding to metabolic reactions and the biomass reaction. Further, there are irreversibility constraints. Altogether, one considers *Nv* = 0 and 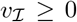, defining the *flux cone* (mathematically, a linear subspace with nonnegativity constraints, that is, an s-cone [19]).

Metabolic pathway analysis aims to identify biologically meaningful routes in a metabolic network, in particular, minimal routes. The (biologically and mathematically) fundamental abstractions of minimal metabolic pathways are *elementary flux modes* (EFMs) [24, 25]. Formally, EFMs are the support-minimal vectors of the flux cone. Most importantly, every element of the flux cone can be written as a conformal sum of EFMs (a sum without cancellations) [19]. In convex analysis, *elementary vectors* (of linear subspaces) have been introduced 25 years before their use in metabolic pathway analysis [23]. Recently, the concept of elementary vectors and the corresponding conformal sum theorems have been extended from linear subspaces to polyhedral cones and polyhedra [19]. Indeed, elementary vectors form *unique* sets of *conformal generators* for linear subspaces, polyhedral cones, and polyhedra.

Given the flux cone, flux balance analysis (FBA) adds linear constraints (e.g. flux bounds) and maximizes (biomass) flux [8]. Altogether, FBA considers a linear program on *Nv* = 0, 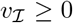, and e.g. *v*^lb^ ≤ *v* ≤ *v*^ub^, defining a *flux polyhedron*. Its elementary vectors are called *elementary flux vectors* (EFVs) [27] and are not support-minimal, in general. As a consequence, maximum flux is attained at EFVs. In other words, EFVs are the missing link between EFM analysis, describing all (stoichiometrically) *feasible* solutions, and FBA, identifying *optimal* solutions [16]. Interestingly, in kinetic models (without flux bounds and with one enzyme capacity constraint), maximum flux is attained at EFMs [20, 29].

Finally, two more classes of elementary vectors have been introduced in metabolic pathway analysis: *elementary conversion modes* (ECMs) of the conversion cone (the projection of the flux cone onto the exchange reactions) [28] and *minimal cut sets* (MCSs) of a dual polyhedron (minimal sets of gene knockouts) [2]. By using ECMs, EFMs need not be enumerated (which is computationally infeasible for genome-scale metabolic models [30]) in order to determine all minimal conversions of substrates into products, and by using MCSs, EFMs need not be enumerated in order to determine all minimal interventions in computational strain design.

Recently, more refined models of cellular growth have been studied, where individual synthesis reactions for macromolecules replace the traditional biomass reaction. In the category of constraint-based models, prominent examples are resource balance analysis (RBA) [11] and metabolism and macromolecular expression (ME) models [17]. In particular, RBA considers linear *capacity constraints*: given the concentration of a catalyst, kinetics implies an upper bound for the flux of the corresponding catalytic reaction. (E.g. for the flux *v* of a metabolic reaction catalyzed by enzyme `E`, one has *v* ≤ *k*^cat^*x*_E_.) In addition, ME models consider the *genotype-phenotype map* of macromolecular synthesis in detail (and the resulting inequality constraints for the fluxes involved). Finally, in the category of kinetic models, whole cell models (also known as self-replicator or self-fabrication models) aim to give a complete picture of cellular processes, however, on a coarse-grained level [5, 18]. In fact, such models often are *hybrid* (kinetic and constraint-based), involving control parameters for optimization and the corresponding constraints.

In this work, we introduce and analyze classes of elementary vectors for *general* models of cellular growth: *Elementary growth modes* (EGMs) are elementary vectors of the *growth cone*; they only depend on stoichiometry and hence apply to general growth models (kinetic or constraint-based). *Elementary growth vectors* are elementary vectors of a *growth polyhedron*; they also depend on growth rate, concentrations, and fluxes and hence apply to constraint-based models. Unlike EFMs (but like ECMs), EGMs are elementary vectors of a general polyhedral cone, and like EFVs, EGVs are elementary vectors of a polyhedron. To demonstrate the relevance of the new concepts, we relate them to the theory of *autocatalytic sets*.

As a case study, we consider whole cell models (simple kinetic models of self-fabrication). To characterize cellular phenotypes with maximum growth rate, we use *elementary growth states* (EGSs) as introduced in [5]. Unlike EGMs, EGSs are not defined for general growth models and also depend on concentrations and growth rate. Still, every EGS can be written as a conformal sum of EGMs. We obtain several new results on EGSs (and hence on growth rate maximization) and simple proofs of existing results.

### Outline and main results

In **Section 2**, we provide new results on elementary vectors in polyhedral geometry, cf. Propositions 1 and 2. In **Section 3**, we introduce *elementary growth modes* and *vectors* for kinetic and constraint-based models of cellular growth, we provide the corresponding conformal sum theorems, cf. Theorems 9, 15 and Corollary 10, we define *minimal autocatalytic sets* of reactions, and we illustrate all new concepts in models of a minimal network, cf. Examples 13 and 16. In **Section 4**, we consider simple kinetic models of self-fabrication and related constraint-based models, we instantiate general definitions and results, and we derive an upper bound for growth rate that only depends on enzyme kinetics (and the set of active enzymes), cf. Theorem 21. In **Section 5**, we study growth rate maximization (via control parameters for ribosome kinetics). In particular, we analyze *elementary growth states* (EGSs) as introduced in [5]. Most importantly, we show that (i) EGSs are support-minimal,

(ii) maximum growth rate is attained at an EGS, and (iii), at every optimal EGS, the ribosome capacity constraint is active, cf. Theorem 28, Proposition 29, Theorem 30, and Corollary 31. Moreover, we find that (E)GSs correspond to (minimal) autocatalytic sets, cf. Proposition 33, and we illustrate the relation between EGSs and EGMs in models of small networks, cf. Examples 34, 35, and 36. Under the assumption of proportional synthesis, we determine the dependence of EGSs on growth rate, cf. Theorem 37, and we study the relation between EGSs and elementary flux modes, cf. Theorems 38 and 39. Finally, in **Section 6**, we discuss the terminology of elementary vectors in metabolic pathway analysis.

In Supplement A, we summarize definitions and results on elementary vectors in polyhedral geometry. In Supplement B, we provide a minimal derivation of the dynamic model of cellular growth. Finally, in Supplement C, we list mathematical issues in [5].

### Mathematical notation

We denote the positive real numbers by ℝ_*>*_ and the nonnegative real numbers by ℝ_≥_. For *x* ∈ ℝ^*n*^, we write *x >* 0 if 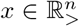, *x* ≥ 0 if 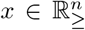, and we denote its *support* by supp(*x*) = {*i* | *x_i_* ≠ 0}. Recall that a nonzero vector *x* ∈ *X* ⊆ ℝ^*n*^ is *support-minimal* if, for all nonzero *x*′ ∈ *X*, supp(*x*′) ⊆ supp(*x*) implies supp(*x*′) = supp(*x*). For *x* ∈ ℝ^*n*^, we define its *sign vector* sign(*x*) ∈ {−, 0, +}^*n*^ by applying the sign function component-wise, that is, sign(*x*)_*i*_ = sign(*x_i_*) for *i* = 1,…, *n*. The relations 0 *<* − and 0 *<* + on {−, 0, +} induce a partial order on {−, 0, +}^*n*^: for *X, Y* ∈ {−, 0, +}^*n*^, we write *X* ≤ *Y* if the inequality holds component-wise. For *x, y* ∈ ℝ^*n*^, we denote the component-wise product by *x* ◦ *y* ∈ ℝ^*n*^, that is, (*x* ◦ *y*)_*i*_ = *x_i_y_i_*. For *n* ∈ N, we write [*n*] = {1*,…, n*}. For *x* ∈ ℝ^*n*^ and index set *I* ⊂ [*n*], we write *x_I_* ∈ ℝ*I* for the corresponding subvector. For *A* ∈ ℝ^*m*×*n*^ and index sets *I* ⊂ [*m*]*, J* ∈ [*n*], we write *A_I,J_* ∈ ℝ^*I*×*J*^ for the corresponding submatrix; if *I* = [*m*], we just write *A*∗*,J* ∈ ℝ^*m*×*J*^.

## 2 Elementary vectors

For the objects of polyhedral geometry (subspaces, cones, polyhedra), there is no *unique* minimal set of generators, in general. However, *elementary vectors* (EVs) form unique sets of *conformal* generators [19, Section 3.4]. For linear subspaces and s-cones (arising from linear subspaces and nonnegativity constraints), elementary vectors are the support-minimal (SM) vectors; for general cones, they are the conformally non-decomposable (cND) vectors; and for general polyhedra, they are the convex-conformally non-decomposable (ccND) vectors plus the cND vectors of the recession cone [19].

In Supplement A, we summarize basic definitions and results for s-cones, general polyhedral cones, and polyhedra; cf. Theorems 43, 44, and 45. Below, we provide new results on special polyhedra (arising from affine subspaces and nonnegativity constraints). In fact, Propositions 1 and 2 will be crucial in the study of elementary growth states; we state them here to complete the mathematical preliminaries before introducing growth models.

### 2.1 Special polyhedra

Recall the definitions of linear and affine subspaces: In a linear subspace *S* ∈ ℝ^*n*^, we have that *x, y* ∈ *S* and *λ, μ* ∈ ℝ imply *λx* + *μy* ∈ *S*. In an affine subspace *T* ∈ ℝ^*n*^, we have that *x, y* ∈ *T* and λ ∈ ℝ imply *λx* + (1 − λ)*y* ∈ *T*. We call an affine subspace *genuine* if it is not also a linear subspace.

#### Proposition 1.

*Let T ⊆ ℝ^*n*^ be a genuine affine subspace. If two SM vectors have the same support, then they are identical*.

*Proof.* Let *e, e*′ ∈ *T* be distinct SM vectors that have the same support. Since *T* is not a linear subspace, they are not scalar multiples. Consider *x* = *λe*+(1 − λ)*e*′ ∈ *T* with λ ∈ ℝ. Indeed, choose λ such that 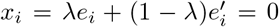 for some *i* ∈ supp(*e*) (with 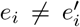). That is, supp(*x*) ⊂ supp(*e*), and *e* is not SM, a contradiction.

The corresponding result for linear subspaces/s-cones is Corollary 42 in Supplement A.1.

#### Proposition 2

*Let T* ⊆ ℝ^*n*^ be a genuine affine subspace and O ⊆ ℝ^n^ be a closed orthant. The vertices of the polyhedron P = *T* ∩ *O are its SM vectors*.

*Proof.* It suffices to show the statement in the notationally simplest case 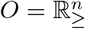. (Then, *x* ∈ *O* is equivalent to *x* ≥ 0.)

(VE ⇒ SM). Assume *x ∈ P* is not SM, that is, there exists *x*^1^ ∈ *P* with supp(*x*^1^) ⊂ supp(*x*), and consider *x*^2^ = *λx* + (1 − λ)*x*^1^ ∈ *T*. Clearly, there exists λ > 1 such that *x*^2^ ≥ 0 (that is, *x*^2^ ∈ *O*) and hence *x*^2^ ∈ *P*. Now, 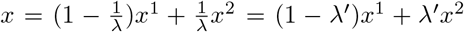 with 0 < λ′ *<* 1, that is, *x* is not a vertex.

(SM ⇒ VE). Assume *x* ∈ *P* is not a vertex, that is, there exist *x*^1^*, x*^2^ ∈ *P* with *x*^1^ ≠ *x*^2^ and 0 *< λ <* 1 such that *x* = *λx*^1^ + (1 − λ)*x*^2^. If supp(*x*^1^) = supp(*x*^2^) = supp(*x*), consider *x*′ = λ′*x*^1^ + (1 − λ′)*x*^2^ ∈ *T*. Clearly, there exists a largest λ′ *>* 1 such that *x*′ ≥ 0 (that is, *x*′ ∈ *O*) and hence *x*′ ∈ *P*. For this λ, supp(*x*′) ⊂ supp(*x*), that is, *x* is not SM.

## 3 Growth models

### Notation

We denote fundamental objects and quantities as follows:

|  |  |  |
| --- | --- | --- |
| Mol | set of molecular species |  |
| Rxn | set of chemical reactions |  |
| $N \in \mathbb{R}^{\text{Mol} \times \text{Rxn}}$ | stoichiometric matrix | (unit: 1) |
| $\rho \in \mathbb{R}_{>}^{\text{Mol}}$ | molar masses | (unit: g/mol) |
| $x \in \mathbb{R}_{\geq}^{\text{Mol}}$ | concentrations | (unit: mol/g) |
| $v(x) \in \mathbb{R}^{\text{Rxn}}$ | reaction rates | (unit: mol/g/s) |

In Supplement B, we provide a minimal derivation of the dynamic model of cellular growth.

A *dynamic* growth model is given by a dynamical system (involving chemical reactions and growth) and a linear constraint representing dry weight (or, alternatively, cell volume):

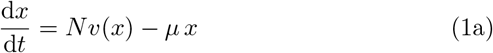

and

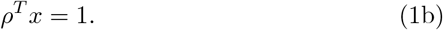

In particular, growth rate is given by

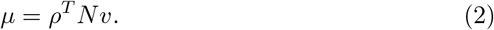

**Note.** At steady state, system (1a) and (2) is equivalent to system (1). However, if (2) is used as a definition of *μ* in (1a), then the mathematical treatment often becomes less transparent.

#### Remark 3

(Conservation laws). In a growth model, there can be no conservation laws. In mathematical terms, ker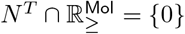

To see this, assume *c^T^ N* = 0 with 0 ≠ *c* ≥ 0, for example, assume *c*_1_ = *c*_2_ = 1 and *c_i_* = 0, otherwise. Then, 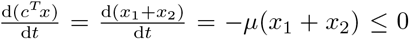, and *μ >* 0 implies *x*_1_ = *x*_2_ = 0 at steady state.

#### Remark 4

(Dependent concentrations). Even if there are no conservation relations, there can be dependent concentrations. (However, biochemically, this is a degenerate case.) In mathematical terms, ker *N^T^* ≠ {0}.

To illustrate this, assume *c^T^ N* = 0 with 0 ≠ *c*, for example, assume *c*_1_ = 1, *c*_2_ = −1, and *c_i_* = 0, otherwise. Then, 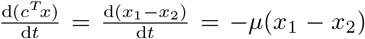, and *μ >* 0 implies *x*_1_ = *x*_2_ at steady state.

In the following, we assume that all dependent variables have been eliminated. In particular, ker *N^T^* = {0}, that is, im *N* = ℝ^Mol^.

#### Elementary vectors

At steady state,

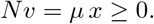

Further, 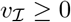, where 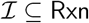 denotes the set of irreversible reactions.

**Note.** In general, all reactions are reversible, but in a given setting, reactions may have a given direction, as determined by thermodynamics.

##### Definition 5

*Growth modes* (GMs) for the dynamic growth model (1) are elements of the *growth cone*

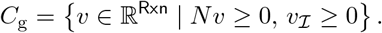

A GM *v C*_g_ has an *associated* growth rate *μ*(*v*) = *ρ^T^ Nv* ≥ 0 and, if *μ*(*v*) *>* 0, an *associated* concentration vector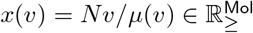

*Elementary* growth modes (EGMs) are conformally non-decomposable GMs.

**Note.** GMs only depend on stoichiometry; in particular, they do not depend on concentrations or growth rate. (But they have an associated growth rate and an associated concentration vector.)

**Note.** The growth cone *C*_g_ is a general polyhedral cone (not an s-cone like the flux cone); its elementary vectors are conformally non-decomposable (but not support-minimal, in general); see e.g. Example 13.

##### Remark 6

(Computation). For the computation of EGMs, the system of inequalities defining *C*_g_ is transformed to a system of equalities by the introduction of slack variables, and SM vectors of the resulting higher-dimensional s-cone are computed by (variants of) the double description method, see e.g. [16].

The definition of GMs immediately implies the scale invariance of associated concentrations.

##### Proposition 7.

*For a GM v* ∈ *C_g_ with associated concentration x*(*v*) *and λ >* 0*, it holds that x*(*λv*) = *x*(*v*).

Further, the assumption ker 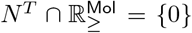 (no conservation laws) allows to characterize zero growth.

##### Proposition 8

*For a GM v* ∈ *C*_g_*, Nv* = 0 *is equivalent to μ*(*v*) = 0.

GMs *v* with *μ*(*v*) = 0 (“zero growth modes”) are flux modes (FMs), that is, elements of the flux cone

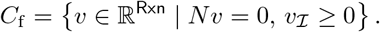

(In general, this is not the flux cone arising from traditional growth models, involving an approximate “biomass reaction”.) EGMs *v* with *μ*(*v*) = 0 are elementary flux modes (EFMs), that is, SM elements of *C*_f_. They are not SM elements of *C*_g_, in general; see e.g. Example 13.

Finally, we apply the general theory of elementary vectors and state the main result of this section.

##### Theorem 9

*Every nonzero GM is a conformal sum of EGMs*.

*Proof.* By Theorem 44 in Supplement A.2: The growth cone *C*_g_ is a general polyhedral cone, and its elementary vectors are the conformally non-decomposable vectors, that is, the EGMs.

In fact, we can be more specific.

##### Corollary 10

*Let v be a nonzero GM with associated growth rate μ*(*v*) =: *μ. Then, there exist (possibly empty) finite sets E*_0_ *and E_μ_ of EGMs with associated growth rates* 0 *and μ, respectively, such that*

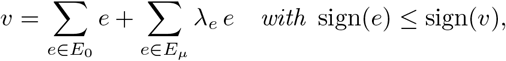

*λ_e_* ≥ 0*, and 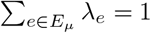. Moreover, if μ >* 0, then 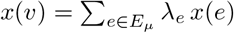.

*Proof.* By Theorem 9, there exist (possibly empty) finite sets *E*_0_ and *E_>_* of EGMs (with associated growth rates 0 and *>*0, respectively) such that

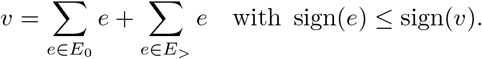

In particular, 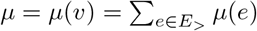. Now

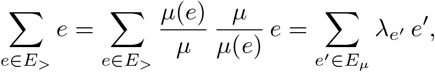

where 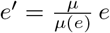 with *μ*(*e*′) = *μ* and 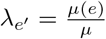 with 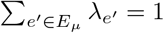. Finally, if *μ >* 0, then 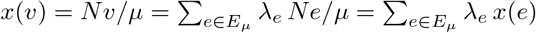.

**Note.** In Corollary 10, we actually fix growth rate which turns the growth cone into a polyhedron. Hence, the result is also an instance of Theorem 45 in Supplement A.3.

#### Autocatalysis

Cellular growth is autocatalytic in the sense that the cell fabricates itself (thereby exchanging substrates/products with the environment). One needs to distinguish this notion of “network autocatalysis” from “autocatalytic subnetworks” (technically: autocatalytic cycles/cores) [3, 4].

A main requirement for network autocatalysis is the existence of a growth mode where *all* species involved in active reactions have nonzero associated concentrations.

##### Definition 11

A GM *v* ∈ *C*_g_ is *strict* if, for every *r* ∈ supp(*v*) ⊆ `Rxn` and *s* ∈ `Mol`, *N_sr_* ≠ 0 implies (*Nv*)_*s*_ > 0.

Clearly, in the overall reaction corresponding to a strict GM, all species appear on the product side, in particular, with a larger stoichiometric coefficient than on the educt side. If a species appears also on the educt side, then it is *formally autocatalytic* (cf. [1]), and one may call the GM itself autocatalytic. In fact, there are several competing notions of autocatalytic species and subnetworks like formally/exclusively autocatalytic and autocatalytic cycles/cores (cf. [1, 3, 4]). Before we state possible definitions for network autocatalysis, we distinguish two modeling approaches.

- **Detailed models** (without individual catalytic reactions) In this approach, catalysis occurs on the level of (small) subnetworks. In particular, individual reactions are not catalytic. For example, a simple catalytic mechanism (involving enzyme `E`, substrate `S`, and product `P`) is given by `E + S ↔ ES ↔ EP ↔ E + P`.
- **Coarse-grained models** (with individual catalytic reactions) In this approach, catalysis occurs on the level of individual reactions. For example, the catalytic mechanism above is written as `E + S ↔ E + P` or 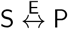. Due to coarse-graining, catalysis cannot be identified from the stoichiometric matrix. Hence, for every catalytic reaction, the corresponding catalyst is specified explicitly.

For **detailed models**, one may call a GM *v* ∈ *C_g_ autocatalytic* if (i) it is strict and (ii) it contains an autocatalytic species or subnetwork. For example, one may require the existence of a formally/exclusively autocatalytic species or a catalytic/autocatalytic subnetwork. Formal definitions and their comparison are subject of future work. In this work, all examples (and the case study in Sections 4 and 5) are specified as coarse-grained models. For **coarse-grained models**, we give a formal definition of network autocatalysis.

##### Definition 12

For coarse-grained models, a GM *v* ∈ *C*_g_ is *basically catalytic* (BC) if there is a catalytic reaction *r* ∈ supp(*v*). Further, a GM *v* ∈ *C*_g_ is *catalytically closed* (CC) if, for every catalytic reaction *r* ∈ supp(*v*), it holds that (*Nv*)_*c*_ > 0 for the corresponding catalyst *c* `Mol`. Finally, a GM *v* ∈ *C_g_* is *autocatalytic* (AC) if it is strict, BC, and CC.

A subset of reactions *S* ⊆ `Rxn` is *autocatalytic* (AC) if there exists an autocatalytic GM *v ∈ C*_g_ with *S* = supp(*v*). A nonempty subset of reactions is *minimally* autocatalytic (MAC) if it is AC and inclusion-minimal.

**Note.** A closure condition is also crucial in classical definitions of “reflexive autocatalysis” [13, 15, 26] and “chemical organizations” [7, 10, 14]. (For detailed models, a closure condition is not required. In that approach, closure is implied by strictness.)

**Note.** One may define an *elementary* autocatalytic GM as an autocatalytic GM with minimal support. However, the definition of autocatalysis goes beyond polyhedral geometry, and hence the term *elementary (vector)* is not appropriate. See also the discussion of terminology in Section 6.

Clearly, network autocatalysis as in Definition 12 implies formal autocatalysis for all catalytic species.

##### Example 13.

Consider the following “minimal network”, involving precursor `P`, “enzyme” `E`, and ribosome `R`:

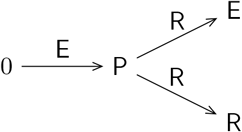

“Metabolism” (the production of the precursor `P`) is catalyzed by the enzyme `E`, and the synthesis of `E` and `R` is catalyzed by the ribosome `R`. The corresponding stoichiometric matrix amounts to

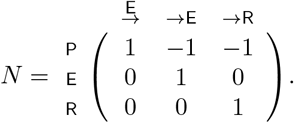

Further, *ρ* = (*ρ*_P_*, ρ*_E_*, ρ*_ℝ_)^*T*^ = *ρ*_P_ (1, 1, 1)^*T*^, by mass conservation. (Normalized) EGMs with associated growth rates and mass fractions are given by

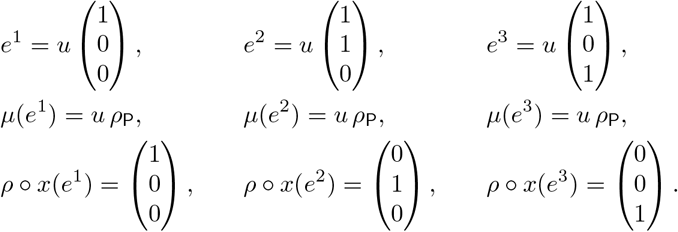

To obtain correct units, we introduced the factor *u* = 1 mol/g/s.

In a (biologically meaningful) kinetic model of the minimal network, all steadystate reaction rates (fluxes) and all (associated) concentrations are nonzero. However, in all EGMs, at least one flux is zero, and only one concentration is nonzero. By definition, EGMs only depend on stoichiometry and hence do not reflect (constraints arising from) kinetics. Still, every flux vector of a kinetic model can be written as a conformal sum of EGMs. See also Examples 16 and 34, where the minimal network is studied in the contexts of constraint-based and kinetic models, respectively.

All EGMs involve catalytic reactions, and hence they are BC. However, for all EGMs, at least one active catalyst has zero (associated) concentration, and hence no EGM is CC. Still, every convex combination *v* = λ_1_ *e*^1^ + λ_2_ *e*^2^ + λ_3_ *e*^3^ of EGMs with λ_1_ ≥ 0, λ_2_*, λ*_3_ *>* 0, λ_1_ + λ_2_ + λ_3_ = 1 (and hence *μ*(*v*) = *u ρ*_P_) has nonzero catalyst concentrations, and hence *v* is CC. If further λ_1_ *>* 0, then also the precursor concentration is nonzero, that is, *v* is strict and ultimately AC. Clearly, *S* = supp(*v*) = `Rxn` (the set of all reactions) is the unique (M)AC set of reactions.

### 3.1 Constraint-based models

For many systems, kinetic models are not yet available, and constraint-based models are used. Steady-state reaction rates (fluxes) *v* are considered as independent variables, that is, the non-linear dependence of the kinetics on the concentrations *x* is neglected. Most importantly, catalytic processes (in the kinetic model) imply linear capacity constraints for *x* and *v*, and additional constraints can be formulated for processes that are not catalytic (in the given model), e.g. lower bounds for concentrations or fluxes. Most compactly, the linear constraints can be can be written as

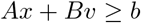

with *A* ∈ ℝ^*m*×Mol^, *B* ∈ ℝ^*m*×Rxn^, and *b* ∈ ℝ^*m*^.

Altogether, *constraint-based* growth models involve steady-state (with irreversibility), dry weight, and additional linear constraints,

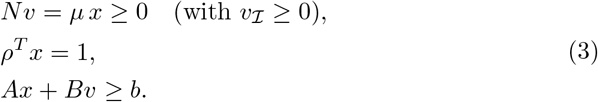

#### Note

For given growth rate *μ* ≥ 0, one may consider the polyhedron

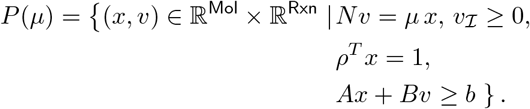

However, for *μ* > 0 and given fluxes *v*, the concentration vector *x* = *Nv/μ* is not an independent variable.

#### Elementary vectors

##### Definition 14.

Let *μ* > 0. *Growth vectors* (GVs) for the constraint-based growth model (3) are elements of the *growth polyhedron*

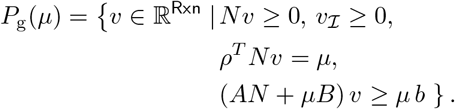

A GV *v* ∈ *P*_g_(*μ*) has an *associated* concentration vector 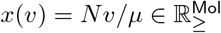.

*Elementary* growth vectors (EGVs) are convex-conformally non-decomposable GVs and conformally non-decomposable elements of the recession cone

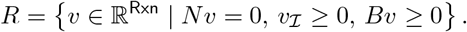

Again, we can apply the general theory of elementary vectors and state our main result.

##### Theorem 15.

*Let v be a GV for growth rate μ* > 0. *Then, there exist finite sets E*_0_ ⊆ *R and E*_*μ*_ ⊆ *P*_g_(*μ*) *of EGVs such that*

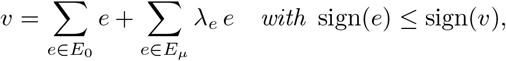

λ_*e*_ ≥ 0, and 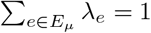. Moreover, 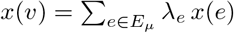.

*Proof.* By Theorem 45 in Supplement A.3: The growth polyhedron *P*_g_(*μ*) is a general polyhedron, and its elementary vectors are the convex-conformally non-decomposable vectors and the conformally non-decomposable vectors of its recession cone.

##### Note.

Every GV is also a GM, and hence every nonzero GV is also a conformal sum of EGMs.

##### Example 16.

Consider the “minimal network” introduced in Example 13 together with the kinetics

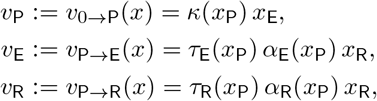

where *α*_E_ + *α*_ℝ_ ≤ 1 due to limited ribosome capacity. The reaction rates are given by particular kinetic functions (depending on the precursor concentration *x*_P_) times the concentrations of the catalyzing molecules (enzyme and ribosome concentrations, *x*_E_ and *x*_ℝ_, respectively).

Now, *κ*(*x*_P_) ≤ *k*^cat^ and 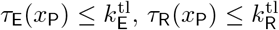, (plus *α*_E_ + *α*_ℝ_ ≤ 1) imply the capacity constraints

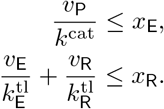

At steady state, *Nv* = *μ x*, that is,

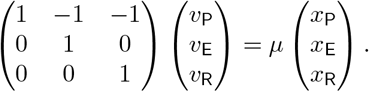

In particular, *x*_E_ = *v*_E_*/μ*, *x*_ℝ_ = *v*_ℝ_*/μ*, and the capacity constraints can be rewritten in terms of fluxes,

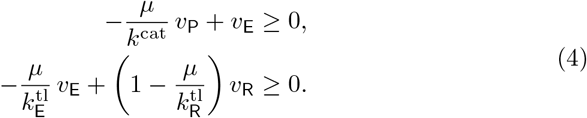

The resulting growth polyhedron amounts to

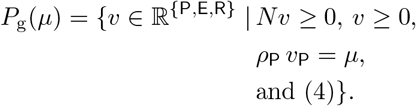

For a quantitative analysis, we set 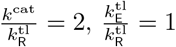, introduce 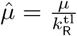, and find 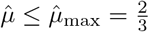. The EGVs and the associated mass fractions are given by

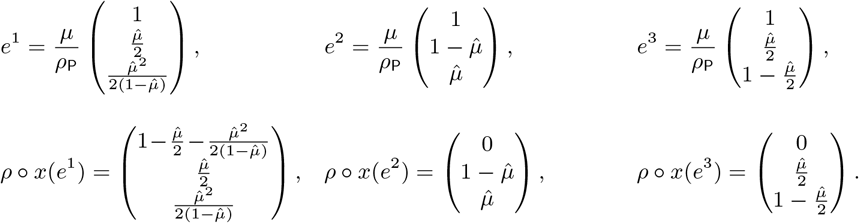

For illustration, the associated mass fractions are shown as functions of growth rate in Figure 1.

**Figure 1:**
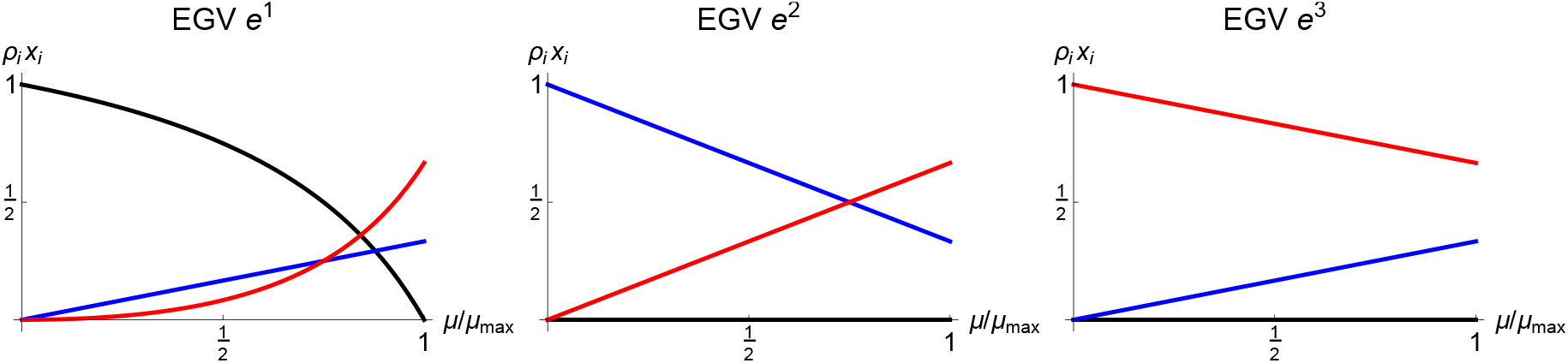
Mass fractions as functions of growth rate for the EGVs *e*^1^, *e*^2^, *e*^3^ of the minimal network. Black/blue/red lines show mass fractions of percursor P/enzyme E/ribosome R.

The defining difference between the EGVs concerns the (in)activity of the inequality constraints *x*_P_ ≥ 0, 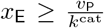, and 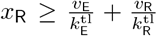. For *μ < μ*_max_, all EGVs have one inactive constraint,

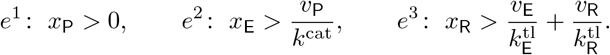

For *μ* = *μ*_max_, all EGVs are identical, *e*^1^ = *e*^2^ = *e*^3^, and all inequality constraints are active, *x*_P_ = 0, 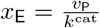, and 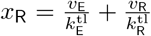. The associated mass fractions are given by 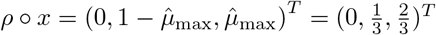.

For all EGVs, all fluxes are nonzero. For *μ < μ*_max_, also all concentrations are nonzero for *e*^1^, whereas *x*_P_ = 0 and *x*_E_*, x*_ℝ_ *>* 0 for *e*^2^ and *e*^3^. Obviously, the capacity constraints do not *fully* reflect kinetics.

All EGVs are BC (since they involve catalytic reactions) and CC (since the active catalysts have nonzero concentrations). If *μ < μ*_max_, then *e*^1^ also has nonzero precursor concentration, that is, *e*^1^ is strict and ultimately AC. As stated in Example 13, *S* = supp(*e*^1^) = `Rxn` is the unique (M)AC set of reactions.

## 4 Simple kinetic models of self-fabrication

Simple kinetic models of self-fabrication as studied in [5, 18] consider metabolism as well as enzyme and ribosome synthesis. In such models, one distinguishes the set of *metabolites* `Met` and the set of (catalytic) *macromolecules* `Mac` (namely, enzymes and the ribosome). Correspondingly, one considers the set of *metabolic reactions* `Rmet` (one reaction per enzyme) and the set of *synthesis reactions* Rsyn (one reaction per macromolecule). To summarize, the sets of molecules `Mol` and reactions `Rxn` are given by

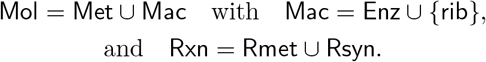

The dynamic growth model (1) for the concentrations 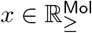 takes the form

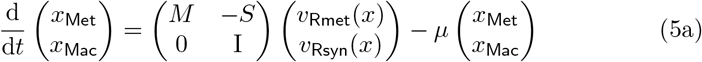

with the dry weight constraint

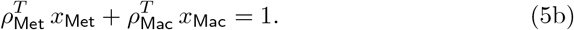

Thereby, we write *M* ∈ ℝ^Met×Rmet^ for the stoichiometric matrix of metabolism (without a “biomass reaction”) and *S* ∈ ℝ^Met×Rsyn^ for the stoichiometric matrix of synthesis, specifying the stoichiometric coefficients of the metabolites (precursors, cofactors,…) in the synthesis reactions (for enzymes and ribosome). Further, we write *v*_Rmet_(*x*) ∈ ℝ^Rmet^ for the rates of the metabolic reactions and 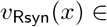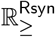 for the synthesis rates; that is, *v*_ℝsyn_(*x*) ≥ 0.

### Remark 17.

Following Remark 4, we assume ker *M*^*T*^ = {0}, that is, im *M* = ℝ^Met^ in the following. Further, we assume that *S* has nonnegative/nonpositive rows, and every column has at least one positive entry.

In simplified notation, the dynamic growth model (5) takes the form:

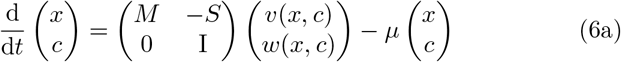

with

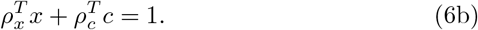

By abuse of the symbols *x* and *v*, we write *x* = *x*_Met_ (for the metabolite concentrations), *v* = *v*_Rmet_ (for the metabolic reactions), and *ρ*_*x*_ = *ρ*_Met_ (for the molar masses of the metabolites). Analogously, we write *c* = *x*_Mac_ (for the enzyme/ribosome concentrations), *w* = *v*_Rsyn_ (for the synthesis reactions), and *ρ_c_* = *ρ*_Mac_ (for the molar masses of enzymes/ribosome).

### Elementary vectors

At steady state,

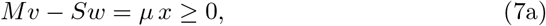

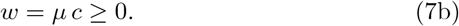

Further, 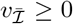, where 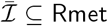 denotes the set of irreversible metabolic reactions. (Implicitly, all synthesis reactions `Rsyn` are assumed to be irreversible.)

In the following, we identify `Rmet` with `Enz` (the metabolic reactions with the catalyzing enzymes) and `Rsyn` with `Mac` (the synthesis reactions with the synthesized macromolecules).

We instantiate Definition 5 and Theorem 9.

#### Definition 18.

*Growth modes* (GMs) for the dynamic growth model (6) are elements of the *growth cone*

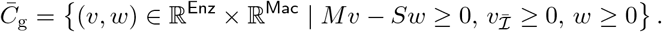

A GM 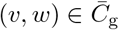 has an *associated* growth rate 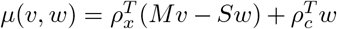 and, if *μ*(*v, w*) > 0, *associated* concentration vectors *x*(*v, w*) = (*Mv − Sw*)*/μ*(*v, w*) and *c*(*v, w*) = *w/μ*(*v, w*).

*Elementary* growth modes (EGMs) are the conformally non-decomposable vectors of 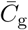.

#### Theorem 19.

*Every nonzero 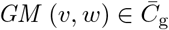 is a conformal sum of EGMs.*

The properties of the matrix *S* imply the following result.

#### Proposition 20.

*Let 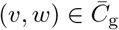 be a GM. If Mv* = 0, *then w* = 0.

Below, we use Theorem 19 and Proposition 20 to derive an upper bound for growth rate that only depends on enzyme kinetics.

### Enzyme kinetics

Often, one considers enzyme kinetics of the form

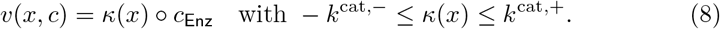

The rate of the metabolic reaction catalyzed by enzyme *i* ∈ `Enz` is given by *v*_*i*_(*x, c*_*i*_) = *κ*_*i*_(*x*) *c*_*i*_, that is, by a particular kinetics times the enzyme concentration.

Enzyme kinetics alone (without specifying ribosome kinetics) implies an upper bound for growth rate (which depends on the set of active enzymes).

#### Theorem 21.

*Assume steady state* (7) *and let enzyme kinetics v*(*x, c*) *be given by* (8). Then, for every x, there is an upper bound for μ, depending on supp(*v*).

*Proof.* By assumption, *Mv − Sw* = *μ x*, *w* = *μ c*, 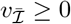, and *v*(*x, c*) = *κ*(*x*) ◦ *c*_Enz_. Obviously, (*v, w*) is a GM. By Theorem 19, (*v, w*) is a conformal sum of EGMs. That is, there exists a set *E* of representative EGMs *e* = (*v*^*e*^, *w*^*e*^) conforming to (*v, w*) (with one representative EGM on each ray of EGMs) such that

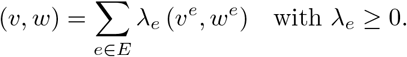

Let *s* = supp(*v*). If *i* ∈ *s*, then *v*_*i*_/κ_*i*_ = *c*_*i*_ = *w*_*i*_/*μ*, that is, *μ* = *w*_*i*_/(*v*_*i*_/*κ*_*i*_), and further

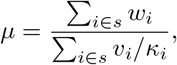

since *r* = *a/b* = *c/d* implies *r* = (*a* + *c*)*/*(*b* + *d*). Using the conformal sum,

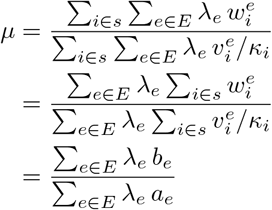

with 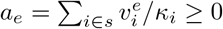 and 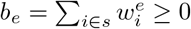.

By Proposition 20, *v*^*e*^ = 0 implies *w*^*e*^ = 0. Equivalently, *a*_*e*_ = 0 implies *b*_*e*_ = 0, that is, *b*_*e*_ > 0 implies *a*_*e*_ > 0. Hence, *μ* as a ratio of linear functions (in the variables λ_*e*_) is bounded. In fact,

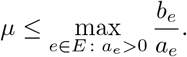

### Ribosome kinetics (with control parameters)

Often, one considers ribosome kinetics of the form

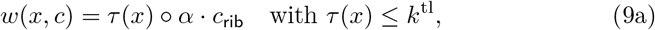

involving the control parameters 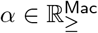 with

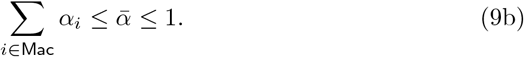

The rate of the synthesis reaction for macromolecule *i* ∈ `Mac` = `Enz` ∪ {`rib`} is given by *w*_*i*_(*x, c*) = *τ*_*i*_(*x*) *α*_*i*_*c*_r_, that is by a particular kinetics times a fraction of the ribosome concentration. Most importantly, the control parameters (ribosome fractions) *α* are used to study growth rate maximization in Section 5.

Steady state (7), enzyme kinetics (8), and ribosome kinetics (9a) imply basic results regarding the case *μ* = 0.

#### Proposition 22.

*Let* 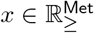 be nonzero. The following statements hold:

- *μ* = 0 ⇒ *w* = 0 ⇒ *Mv* = 0.
- *c*_rib_ = 0 ⇒ *w* = 0 ⇒ *μ* = 0 ∨ *c*_Enz_ = 0. *c*_Enz_ = 0 ⇒ *v* = 0 ∧ *w*_Enz_ = 0 ⇒ *w*_rib_ = 0 ⇒ *μ* = 0. Hence, *c*_rib_ = 0 ⇒ *μ* = 0.

Conversely, *μ* = 0 ⇒ *c*_rib_ = 0 does not hold if *τ* (*x*) ◦; *α* = 0 (a degenerate case). In the following, we often assume *μ* > 0 (and hence *c*_rib_, *w*_rib_, *τ*_rib_, *α*_rib_ > 0).

### 4.1 Related constraint-based models

In the rest of this section, we specify a constraint-based model corresponding to the simple kinetic model of self-fabrication.

First, enzyme kinetics (8) implies the capacity constraints

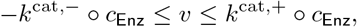

which can be written as

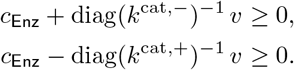

Second, ribosome kinetics (9a) and the capacity constraint (9b) imply

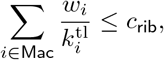

that is,

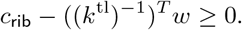

Altogether,

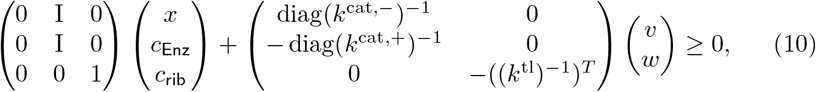

which is of the general form 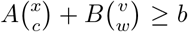. Here, the metabolite concentrations *x* do not contribute to the constraints, and the right-hand side *b* is zero.

In general, constraint-based growth models (3) involve steady state (with irreversibility), dry weight, and additional linear constraints. Here, the model involves steady state (7) (with irreversibility 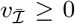), dry weight (6b), and the additional linear constraints (10).

After eliminating enzyme/ribosome concentrations *c* via steady state (7b), that is, *c* = *w/μ*, the additional constraints only involve the reaction rates (fluxes) *v* and *w*,

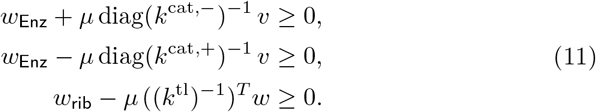

By Definition 14, the growth polyhedron for the constraint-based model is given by

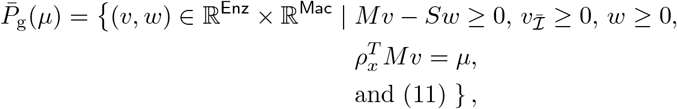

and GVs and EGVs are defined accordingly.

## 5 Growth rate maximization

We consider the simple kinetic model of self-fabrication and study the problem of growth rate maximization. In fact, we maximize growth rate at steady state (7) and for given enzyme kinetics (8), ribosome kinetics (9a) with ribosome capacity (9b), and dry weight (6b).

Thereby, we further simplify notation and introduce the abbreviations `E` = `Enz` and `r` = `rib` and hence `Mac` = `E` ∪ {`r`}.

### Problem 23.

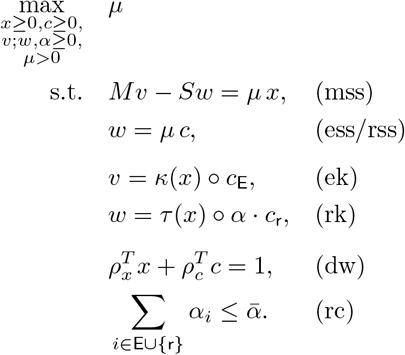

### Legend.

mss/ess/rss = metabolite/enzyme/ribosome steady state, ek/rk = enzyme/ribosome kinetics, dw = dry weight, rc = ribosome capacity

### Remark 24.

We write *x* ≥ 0 in the optimization problem. In fact, we assume *x* ∈ *D*, where *D* is a compact (bounded and closed) set. As a consequence, the maximum is attained.

In the following, we eliminate the enzyme/ribosome concentrations *c* and the fluxes *v, w* as independent optimization variables. That is, we only keep the control parameters (ribosome fractions) *α* as well as the metabolite concentrations *x* and growth rate *μ*.

Enzyme/ribosome steady state (7b) and ribosome kinetics (9a) imply

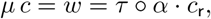

that is,

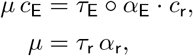

which yields *c*_E_ in terms of *α*_E_ (and *x*, *μ*).

Now, dry weight (6b) implies

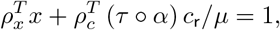

that is,

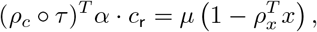

which yields *c*_r_ in terms of *α* (and *x*, *μ*).

As a consequence, all enzyme concentrations, all fluxes, and also growth rate are multiples of the ribosome concentration *c*_r_,

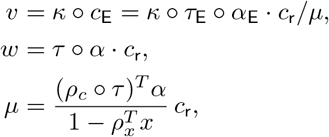

and we divide metabolite steady state by *c*_r_ > 0. Altogether, we formulate growth rate maximization in terms of *α* (and *x*, *μ*).

### Problem 25

(in terms of *α*).

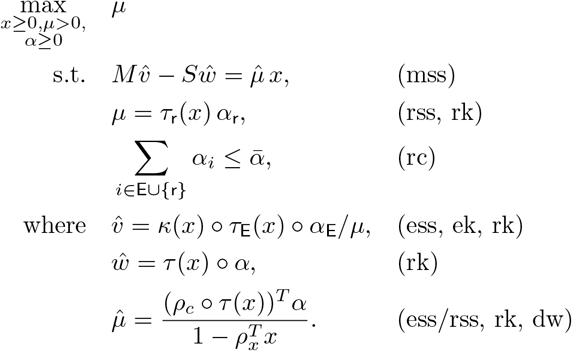

### Legend.

Labels denote the dependence of the equations/inequalities on the original constraints in Problem 23.

### Note.

The last three equations are not constraints, but definitions of 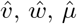 in terms of *α* (and *x*, *μ*).

### Elementary vectors

We follow the definition in [5], but write it in a more transparent way which allows to recognize the origin of the constraints and the resulting affine subspace at the same time.

#### Definition 26.

Let *μ* > 0. *Growth states* (GSs) for Problem 25 are elements of the polyhedron

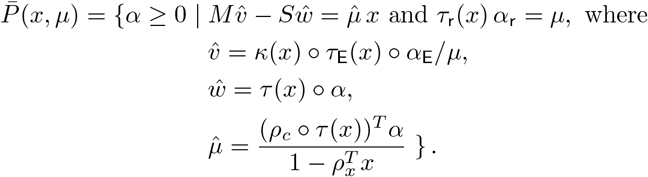

*Elementary* growth states (EGSs) are the vertices of 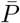.

#### Note.

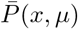 is the intersection of an affine subspace with the nonnegative orthant and hence unbounded, in general; that is, 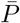 is a polyhedron; see also Example 35. The elementary vectors of a polyhedron are its convex-conformally non-decomposable (ccND) vectors and the conformally non-decomposable vectors of its recession cone; see Supplement A.3. We define EGSs as ccND vectors which agree with the vertices due to nonnegativity.

By the definition of EGSs (as vertices of a polyhedron), we have the following (necessarily conformal) sum theorem.

#### Proposition 27.

*For given polyhedron 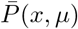, every convex combination of EGSs is a GS, but not vice versa, in general.*

By Theorem 21, for every *x*, there is an upper bound for *μ*, depending on supp(*v*). (And ribosome capacity further limits growth rate.) Clearly, 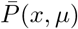 is empty for *μ* above the global upper bound. Moreover, different EGSs have different upper bounds, in general, and the number of vertices of 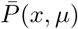 depends on *μ*; see also Example 36.

The following result is stated (with an incorrect proof) in [5]. We give a simple proof which uses a polyhedral geometry argument and does not require the implicit function theorem (IFT).

#### Theorem 28.

The maximum of Problem 25 is attained at an EGS.

*Proof.* Let 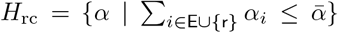 be the halfspace given by ribosome capacity. For given *x* and *μ*, in particular, for their optimal values, the set of feasible *α* is given by the polytope 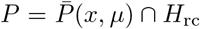. If *P* ≠ ϕ (as for optimal *x* and *μ*), then at least one vertex of 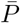 is a vertex of 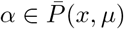, that is, an EGS.

#### Note.

Let *x* and *μ* be optimal. By the proof above, the maximum is attained at *every* EGS 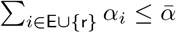 that fulfills 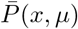.

Next, we show that EGSs (vertices of 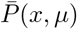) are support-minimal, a crucial result that is missing in [5].

#### Proposition 29.

*EGSs are the support-minimal vectors of the polyhedron* 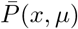.

*Proof.* The polyhedron 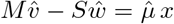 is given by the linear equations 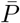 (homogeneous in *α*), the inhomogeneous equation *τ*_r_(*x*) *α*_r_ = *μ*, and nonnegativity *α* ≥ 0, that is, it is the intersection of a genuine affine subspace with the nonnegative orthant. By Proposition 2, the vertices of 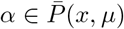 are its support-minimal vectors.

Using Proposition 29, we show that EGSs with the same support also exist in a neighbourhood (of *x* and *μ*).

#### Theorem 30.

*Let* 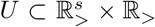 *be an EGS with s* = supp(*x*)*. Then, there exists an open neighborhood* 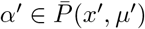 *of* (*x_s_, μ*) *such that, for all* (*ξ, μ*′) ∈ *U, α can be continuously extended to an EGS* 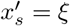 *with* supp(*x*′) = *s*, 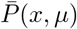, *and* supp(*α*′) = supp(*α*).

*Proof.* Altogether, the polyhedron 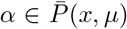 is given by inhomogeneous linear equations

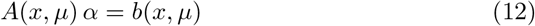

and *α* ≥ 0, that is, it is the intersection of a genuine affine subspace with the non-negative orthant. By Proposition 29, the EGS 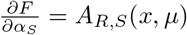 is support-minimal. By Proposition 1, *α* is the unique solution of (12) with support *S* = supp(*α*). Hence, there is an invertible square submatrix *A_R,S_* (*x, μ*) of *A* (with |*R*| = |*S*|) and a subvector *b_R_*(*x, μ*) of *b* such that the subvector *α_S_* > 0 of *α* fulfills

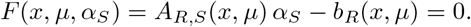

Clearly, 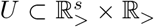 is invertible.

Now, consider *x_s_*, the nonzero part of *x*. By the IFT, there exists an open neighborhood 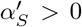 of (*x_s_, μ*) such that, for all (*ξ, μ*′) *U*, *α_S_* can be continuously extended to 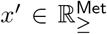. With 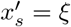 given by supp(*x*′) = *s* and 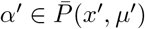, the EGS *α* can be continuously extended to the EGS 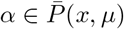 with supp(*α*′) = *S*.

A weaker (and incorrect) version of Theorem 30 is stated in [5]. In fact, it has an unnecessary assumption on the support of *α*, and it lacks an assumption on the support of *x*.

Finally, using Theorem 30, we can further characterize optimal solutions (of growth rate maximization).

#### Corollary 31.

*At every optimal EGS of Problem 25, the ribosome capacity constraint is active*.

*Proof.* Let 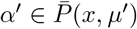 be an EGS for maximum *μ* (and corresponding optimal *x*). By Theorem 30, there is *μ*′ > *μ* such that *α* can be continuously extended to an EGS 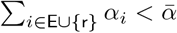 with supp(*α*′) = supp(*α*). Now, assume 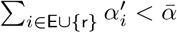. By continuity (of *α* as a function of *μ*), also 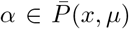. Hence, (*x, μ*′, *α*′) is a feasible solution of Problem 25 which contradicts the maximality of *μ*.

In the main text of [5], it is (incorrectly) claimed that, at maximum growth rate, the ribosome capacity constraint need not be active (but some ribosome fraction may become zero).

### 5.1 EGSs and minimal autocatalytic sets

By Definitions 26 and 18, every GS has a corresponding GM.

#### Proposition 32.

*Let* 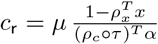 *be a GS, and let v* = *κ* ◦; *τ*_E_ ◦; *α*_E_ · *c*_R_*/μ and w* = *τ* ◦; *α* · *c*_R_ *with* 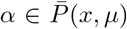 *be the corresponding (metabolic and synthesis) fluxes. Then,* (*v, w*) *is a GM with associated growth rate μ*(*v, w*) = *μ.*

Moreover, by Definition 12, the GM corresponding to a GS is autocatalytic. Thereby, we make the (biochemically meaningful) assumption on (enzyme and ribosome) kinetics that, if a species *s* is consumed in a reaction *r* (short: *s* → 0), then *v_r_* > 0 implies *x_s_* > 0.

#### Proposition 33.

*Let* 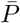 *be a GS, and let* (*v, w*) *be the corresponding GM. Then,* (*v, w*) *is AC, and hence* supp(*v, w*) *is an AC subset of reactions*.

*Proof.* First, we show that (*v, w*) is strict. Assume that *i* ∈ supp(*v*) ⊆ `E` and *M_si_* ≠ 0 for some species *s* ∈ `Met`. The terms *M_si′_ v_i′_* and −*S_sj′_ w_j′_* in the sum (*Mv* − *Sw*)_*s*_ ≥ 0 can be positive, negative, or zero, and there is a positive term. If there is a negative term, then there is a reaction that consumes *s* and hence *x_s_* > 0. In any case, (*Mv* − *Sw*)_*s*_ = *μx_s_* > 0. Alternatively, assume that *j* ∈ supp(*w*) ⊆ `E` ∪ {`r`}. Regarding species *s* ∈ `Met`, the proof is analogous. Regarding species *s* ∈ `E` ∪ {`r`}, obviously I_*sj*_ /= 0 if and only if *s* = *j* and hence (I *w*)_*s*_ = *w_j_* > 0.

Since all reactions are catalytic, (*v, w*) is BC. It remains to show that (*v, w*) is CC. Assume that *i* ∈ supp(*v*) ⊆ `E`. Then, *τ_i_, α_i_, c*_R_ > 0 and hence (I *w*)_*i*_ = *w_i_* > 0. Finally, *τ*_R_, *α*_R_ > 0 (by the definition of 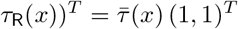) and hence (I *w*)_R_ = *w*_R_ > 0.

We have shown that GSs correspond to autocatalytic (AC) sets of reactions. One may conjecture that EGSs correspond to minimal autocatalytic (MAC) sets of reactions. However, the formalization of this statement is beyond the scope of this work. Recall that the definition of EGSs depends on enzyme and ribosome kinetics (with control parameters), that is, ultimately on metabolite concentrations, whereas the definition of MAC sets of reactions only depends on stoichiometry.

### 5.2 EGSs and EGMs

In three examples, we determine the EGSs and write them as conformal sums of EGMs.

First, we consider the minimal network studied in Examples 13 and 16, and then we extend it in two different ways: we assume that “metabolism” consists of (i) two anti-parallel pathways or (ii) two parallel (alternative) pathways. Thereby, we demonstrate two properties of the polyhedron of growth states (GSs): (i) it can be unbounded, and (ii) the number of its vertices (EGSs) can change with growth rate.

As in Examples 13 and 16, we use E to denote an individual enzyme (whereas `E` = `Enz` is used as an abbreviation for the set of all enzymes in all other parts of Section 5) and `R` to denote the ribosome (instead of `r` = `rib`).

#### Example 34.

Consider the minimal network studied in Examples 13 and 16:

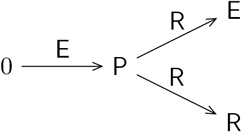

The corresponding matrices of metabolism and synthesis amount to

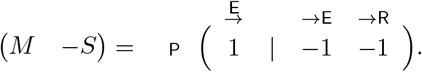

Further, *ρ_x_* = *ρ*_P_ =: *ρ* and *ρ_c_* = (*ρ*_E_, *ρ*_R_)^*T*^ = *ρ* (1, 1)^*T*^, by mass conservation. Analogously, we assume *τ* (*x*) = (*τ*_E_(*x*), 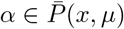.

(Normalized) EGMs are given by

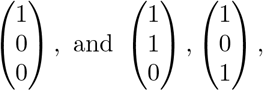

multiplied by a factor *u* = 1 mol/g/s. All EGMs have associated growth rate *μ* = *u ρ*. By Definition 18, the first EGM has associated concentrations *x* = *x*_P_ = 1 and *c* = (*c*_E_, *c*_R_)^*T*^ = 0, and the other two have *x* = 0 and *c* /= 0.

Let 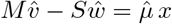 be a GS, and let

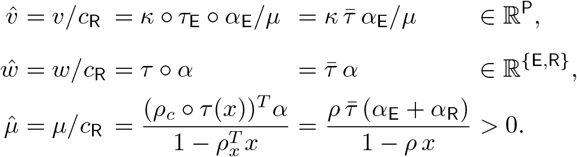

By Definition 26, 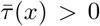 and *τ*_R_ *α*_R_ = *μ*. Explicitly,

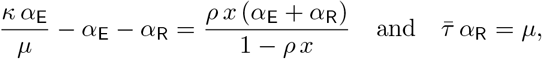

which has the unique solution

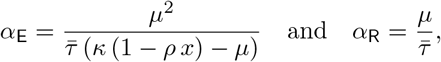

thereby assuming *κ*(*x*), 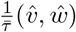. That is, in the minimal network, there is a unique GS and hence a unique EGS. Moreover,

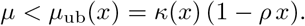

In fact, the ribosome capacity constraint *α*_E_ + *α*_R_ ≤ 1 (which is not part of Definition 26) further limits growth rate,

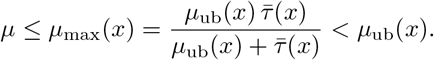

By Proposition 32, the dimensionless vector 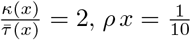 is the corresponding (scaled) GM, and further, by Theorem 19, it is a conformal sum of (scaled) EGMs,

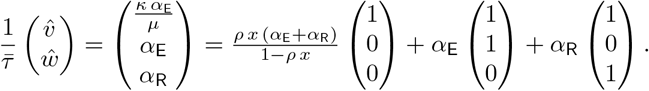

The corresponding mass fractions are given by

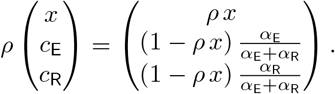

For a quantitative analysis, we fix *x* = *x*_P_; in particular, we set 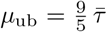 and find 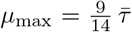 and 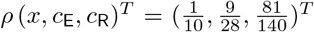. The resulting mass fractions are shown as functions of growth rate in Figure 2. Interestingly, they are linear. For *μ* = *μ*_max_, 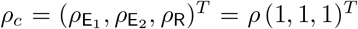. Compare with Figure 1 and recall that EGVs do not depend on the precursor concentration *x*_P_, but have an associated *x*_P_.

**Figure 2:**
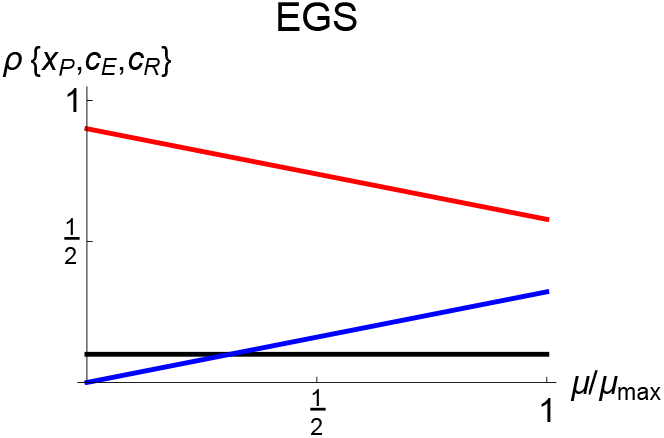
Mass fractions as functions of growth rate for the unique EGS of the minimal network (for fixed precursor concentration). Black/blue/red lines show mass fractions of percursor P/enzyme E/ribosome R.

By Proposition 33, the GM (*v, w*) corresponding to the unique (E)GS is AC. As stated in Example 13, *S* = supp(*v, w*) = Rxn is the unique (M)AC set of reactions.

#### Example 35.

“Metabolism” (the production and consumption of the precursor `P`) is catalyzed by the “enzymes” `E`_1_ and `E`_2_, respectively, and the synthesis of `E`_1_, `E`_2_, and `R` is catalyzed by the ribosome `R`:

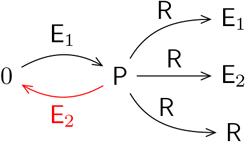

The corresponding matrices of metabolism and synthesis amount to

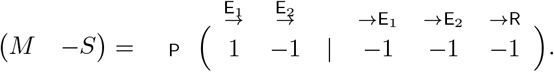

Further, *ρ_x_* = *ρ*_P_ =: *ρ* and 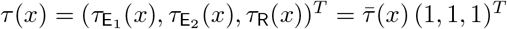, by mass conservation. Analogously, we assume 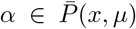.

(Normalized) EGMs are given by

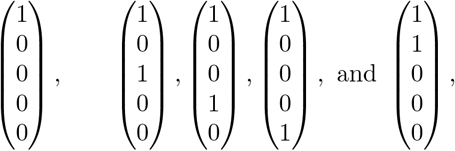

multiplied by a factor *u* = 1 mol/g/s. All EGMs except the last one have associated growth rate *μ* = *u ρ*, and the last one has *μ* = 0. By Definition 18, the first EGM has associated concentrations *x* = 1 and *c* = 0, and the second group of EGMs has *x* = 0 and *c* ≠ 0.

There is only one EGS 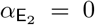, and it has 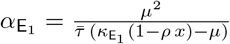. This reduces the problem to the minimal network (with E = E_1_), and the unique solution is given by 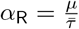 and 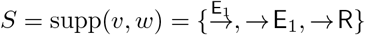.

By Proposition 33, the GM (*v, w*) corresponding to the unique EGS is AC. In fact, 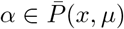 is the unique MAC set of reactions.

In general, a GS 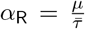 is given by

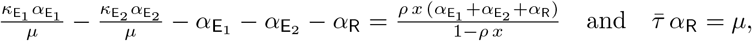

and the corresponding (scaled) GM can be specified as a conformal sum of (scaled) EGMs,

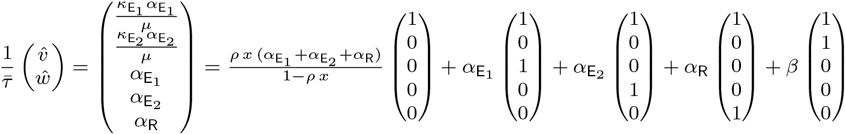

with 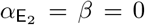 and *β* ≥ 0. Clearly, for the unique EGS, 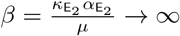, and for unbounded GSs, 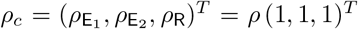.

#### Example 36 Alternative pathways

“Metabolism” (the production of the precursor `P`) is catalyzed alternatively by the “enzymes” `E`_1_ and `E`_2_, and the synthesis of `E`_1_, `E`_2_, and `R` is catalyzed by the ribosome `R`:

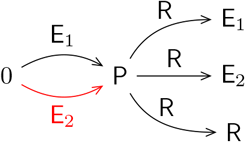

The corresponding matrices of metabolism and synthesis amount to

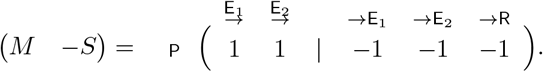

Further, *ρ_x_* = *ρ*_P_ =: *ρ* and 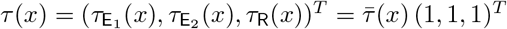, by mass conservation. Analogously, we assume 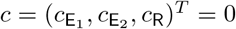.

(Normalized) EGMs are given by

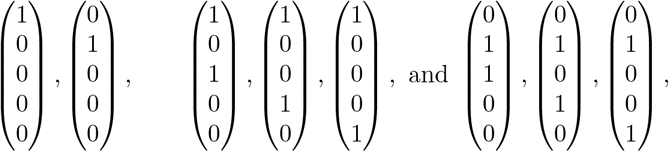

multiplied by a factor *u* = 1 mol/g/s. All EGMs have associated growth rate *μ* = *u ρ*. By Definition 18, the first group has associated concentrations *x* = *x*_P_ = 1 and 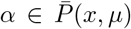 and the other two groups have *x* = 0 and *c* ≠ 0.

Let 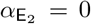 be the EGS with 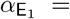. This reduces the problem to the minimal network (with E = E_1_), and the unique solution is given by 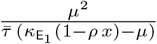 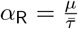 and 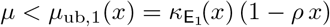. Moreover, 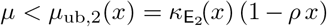.

Analogously, 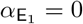 for the EGS *α* with 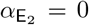. Hence, depending on *μ* (and *x*), there are either two, one, or zero EGSs.

By Proposition 33, the GMs (*v, w*) corresponding to the EGS *α* with 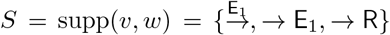 is AC. In fact, 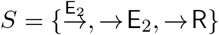 is a MAC set of reactions. Analogously, 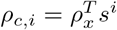 is a MAC set of reactions.

### Proportional synthesis

The final results of this work concern the dependence of EGSs on growth rate and the relation between EGSs and EFMs.

For simplicity, we assume that the reaction vectors of the synthesis reactions (for enzymes and ribosome) are proportional. Let *s^i^* ∈ ℝ^Met^ be the column of *S* ∈ ℝ^Met×(E∪{r})^ corresponding to *i* ∈ `E` ∪ {`r`} (and note that 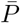). Now, let *i*\* ∈ `E` be a representative enzyme with corresponding column *s*:= *s^i*^* and molar mass *ρ*:= *ρ_c,i*_*. Then,

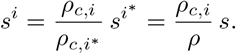

In matrix notation,

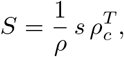

involving a dyadic product. At steady state (7),

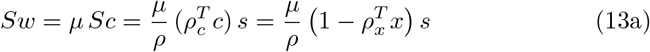

and further

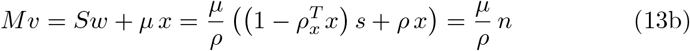

with the dimensionless vector

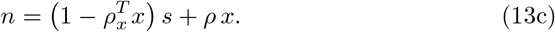

#### Note.

The vector *n* summarizes the effects of both synthesis reactions and dilution by growth.

#### 5.3 EGSs and growth rate

Under the assumption of proportional synthesis, growth states (GSs) are elements of the (simplified) polyhedron

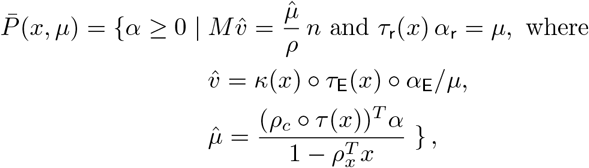

and elementary growth states (EGSs) are the vertices of 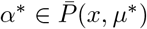. For fixed *x*, the dependence of EGSs on *μ* can be stated explicitly.

##### Theorem 37.

*Assume proportional synthesis, and let* 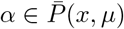 *be an EGS. Then, for every EGS* 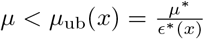 *with* supp(*α*) = supp(*α*\*), *it holds that*

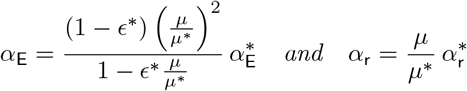

*with*

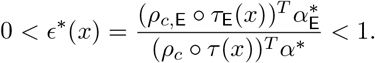

*In particular*, 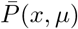. *The aggregated mass fractions of metabolites, enzymes, and ribosome are given by*

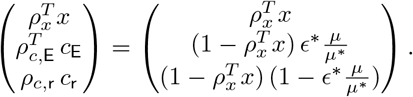

*Proof.* The definition of 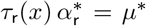 implies 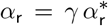, *τ*_R_(*x*) *α*_R_ = *μ*, and hence 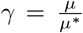 with 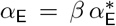. Analogously, assume 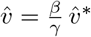 with *β* > 0. Then,

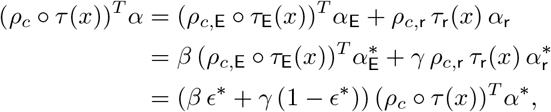

and hence also 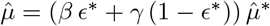. Now, consider 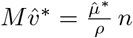 and 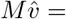 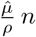. In the transition from *μ*\* to *μ*, the left- and right-hand sides scale with 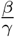 and *β ϵ*\* + *γ* (1 − *ϵ*\*), respectively. Hence, the assumption above is consistent if 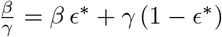, that is, 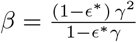.

To determine the mass fractions, observe that

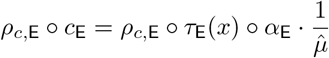

scales with 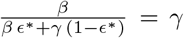. Clearly, 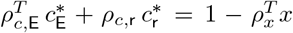, and finally, 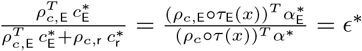.

##### Note.

All control parameters (ribosome fractions) scale with the same nonlinear factor (as a function of growth rate), whereas ribosome synthesis scales linearly. Interestingly, enzyme mass fractions scale linearly (and hence also ribosome mass fraction). See also Figure 2 for the EGS of the minimal network.

#### 5.4 EGSs and EFMs

Under the assumption of proportional synthesis, we introduce a “biomass reaction” `bm` with reaction vector *n*_bm_ = −*n* and reaction rate 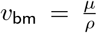 (unit: mol/g/s) corresponding to growth rate *μ* (unit: 1/s).

##### Note.

For the definition of reaction rate, a scaling factor (unit: g/mol) is required; for simplicity, we choose the molar mass *ρ* of the representative protein.

At steady state, 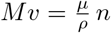, cf. Equation (13b), that is,

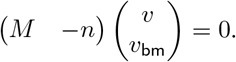

Indeed, *M* −*n* ℝ^Met×(E∪{bm})^ is the stoichiometric matrix of traditional growth models, that is, the stoichiometric matrix of metabolism extended by the biomass reaction vector.

Flux modes (FMs) are elements of the corresponding flux cone

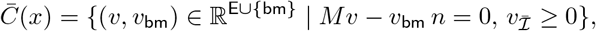

and elementary flux modes (EFMs) are the support-minimal vectors of 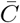. The flux cone is an s-cone, and by Theorem 43 in Supplement A.1, every FM is a conformal sum of EFMs.

We characterize when the FM corresponding to an EGS is an EFM itself.

##### Theorem 38.

*Assume proportional synthesis, and let* 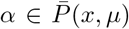 *be an EGS with s* = supp(*α*_E_). *Further, let v* ∈ ℝ^E^ *be the corresponding metabolic fluxes and v*_bm_ = *μ/ρ. Then*, 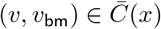 *is an EFM if and only if M*_∗,*s*_ *has rank* |*s*|.

*Proof.* By assumption, *M*_*,*s*_*v_s_* −*v*_bm_(*n* = 0. By)Proposition 41 in Supplement A.1, (*v*, *v*_bm_) is an EFM if and only if ker (*M*_*,*s*_ −*n*) is one-dimensional. Equivalently, ker *M*_∗,*s*_ = {0}, that is, *M*_∗,*s*_ has rank |*s*|.

Alternatively, we introduce a biomass reaction that considers only the synthesis reactions, but not dilution by growth. Indeed, 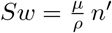 with 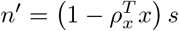, cf. Equation (13a), and we define 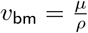 and *n*_bm_ = −*n*′.

At steady state, *Mv* − *Sw* = *μ x*, cf. Equation (7), that is,

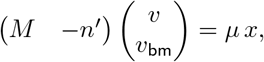

and the corresponding flux cone is given by

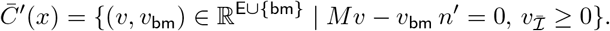

In traditional growth models, this definition of the flux cone arises from the assumption *Mv* ≈ *Sw*, that is, *μx* ≪ |*Mv*|, |*Sw*| (with absolute values taken component-wise). This assumption has been made in FBA models, but is not always justified (in particular, if (*Sw*)_*i*_ = 0 and hence (*Mv*)_*i*_ = *μ x_i_*).

Again, we characterize when the FM corresponding to an EGS is an EFM itself.

##### Theorem 39.

*Assume proportional synthesis, and let* 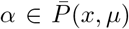 *be an EGS with s* = supp(*α*_E_). *Further, let v* ∈ ℝE *be the corresponding metabolic fluxes and v*_bm_ = *μ/ρ. Then, the following statements are equivalent*:

i. *v* = *v*^1^ + *v*^2^, *where* 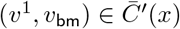 *is an EFM and Mv*^2^ = *μ x*.
ii. *M*_*,*s*_ *has rank* |*s*|. *Proof.* See Theorem 38.

##### Remark 40.

In Theorem 39, both fluxes are proportional to growth rate, 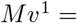 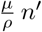 and *Mv*^2^ = *μ x*. If the contribution of metabolites to dry weight is negligible, then the vector *v*^2^ can be seen as a perturbation of the EFM *v*^1^. Indeed, 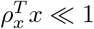 implies 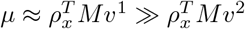.

## 6 Terminology

*Elementary flux modes* (EFMs) are defined as support-minimal vectors of the flux cone, for general (growth or non-growth) models, not just for traditional growth models (involving an approximative biomass reaction). In particular, EFMs only depend on stoichiometry. Similarly, elementary flux *vectors* (EFVs) are defined for general models with additional linear constraints.

In this work, we consider general *growth* models, and we define *elementary growth modes* (EGMs) as conformally non-decomposable vectors of the growth cone. Importantly, EGMs only depend on stoichiometry. Similarly, we introduce elementary growth *vectors* (EGVs) for general growth models with additional linear constraints (such as FBA and ME models).

For simple kinetic models of self-fabrication (with control parameters), *elementary growth states* (EGSs) were introduced as vertices of a polyhedron (in the space of ribosome fractions) in [5]. In this work, we have shown that EGSs are support-minimal. Hence, it makes sense to give a name to the supports of EGSs. Unfortunately, these sets were called “elementary growth modes (EGMs)” in [5], without defining “growth modes (GMs)”. For several reasons, this terminology is not appropriate: Unlike EFMs, (i) “EGMs” are not defined for general growth models, but just for simple kinetic models of self-fabrication, (ii) “EGMs” do not depend on stoichiometry only, but also on growth rate and concentrations, in particular, on enzyme and ribosome kinetics, and (iii) “EGMs” are not elementary (vectors) in the sense of polyhedral geometry, but supports (sets). In this work, we have shown that supports of (E)GSs correspond to *(minimal) autocatalytic sets* of reactions, defined for coarse-grained (but otherwise general) growth models. In a follow-up paper, we elaborate on the relation between detailed growth models and autocatalysis.

## Acknowledgements

We thank Ralf Steuer for discussions on kinetic/constraint-based models of cellular growth (see also the review [21]) and Christoph Flamm for discussions on autocatalytic sets. SM was supported by the Austrian Science Fund (FWF), project P33218.

## Supplementary material

### A Elementary vectors

Below, we summarize basic definitions and results for s-cones, general polyhedral cones, and polyhedra.

#### A.1 S-cones

Given a linear subspace *S* ⊆ ℝ^*n*^ and an index set *I* ⊆ [*n*], an *s-cone* (special cone, subspace cone) is given by *C*(*S, I*) = {*x* ∈ ℝ^*n*^ | *x* ∈ *S, x_I_* ≥ 0}. Note that a linear subspace is an s-cone, *S* = *C*(*S,* ∅).

Before we state the fundamental result for s-cones, we give an alternative definition of SM vectors, in order to demonstrate the underlying proof techniques.

##### Proposition 41.

*Let C*(*S, I*) *be an s-cone. A nonzero vector x* ∈ *C*(*S, I*) *with s* = supp(*x*) *is SM if and only if the linear subspace S_s_* = *x S* supp(*x*) *s has dimension one*.

*Proof.* Obviously, *x* ∈ *S_s_*. (⇐) If dim *S_s_* = 1, then every element of *S_s_* is a scalar multiple of *x*, and hence *x* is SM. (⇒) If *x* is SM, assume dim *S_s_* > 1. Then, there exists *x*′ ∈ *S_s_* (not necessarily *x*′ ∈ *C*(*S, I*)) which is not a scalar multiple of *x*. Consider *x*\* = *x* − *λx*′ with λ ∈ ℝ. Note that *x*\* ≠ 0. Indeed, choose λ such that sign(*x*\*) ≤ sign(*x*) (no sign change of *x*\* relative to *x*) and 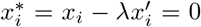 for some *i* ∈ *S_s_*. Then 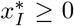, and supp(*x*\*) ⊂ supp(*x*). To summarize, 0 ≠ *x*\* ∈ *C*(*S, I*), and *x* is not SM, a contradiction.

##### Corollary 42.

*Let C*(*S, I*) *be an s-cone. If two SM vectors have the same support, then they are scalar multiples.*

A vector *x ∈ C*(*S, I*) is *elementary* if it is SM. (For linear subspaces, the definition of elementary vectors (EVs) as SM vectors was given in [23].)

The following result is fundamental. See [19, Theorem 3] based on [23, Theorem 1].

##### Theorem 43.

*Let C*(*S, I*) *be an s-cone. Every nonzero vector x ∈ C*(*S, I*) *is a conformal sum of EVs. That is, there exists a finite set E of EVs such that*

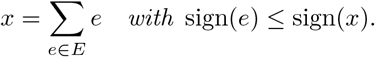

*The set E can be chosen such that* |*E*| ≤ dim(*S*) *and* |*E*| ≤ | supp(*x*)|.

#### A.2 General polyhedral cones

Let *C* be a polyhedral cone, that is, *C* = {*x* ∈ ℝ^*n*^ | *Ax* ≥ 0} for some *A* ∈ ℝ*m*×*n*. A nonzero vector *x* ∈ *C* is *conformally non-decomposable* (cND) if, for all nonzero *x*^1^, *x*^2^ ∈ *C* with sign(*x*^1^), sign(*x*^2^) ≤ sign(*x*), the decomposition *x* = *x*^1^ + *x*^2^ implies *x*^1^ = *λx*^2^ with λ > 0. A vector *x* ∈ *C* is *elementary* if it is cND.

By defining EVs as cND vectors (instead of SM vectors), Theorem 43 can be extended to general polyhedral cones. See [19, Theorem 8].

##### Theorem 44.

*Let C* = {*x* ℝ^*n*^ ≥ A*x* 0 *be a polyhedral cone. Every nonzero vector x C is a conformal sum of EVs. That is, there exists a finite set E of EVs such that*

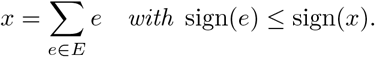

*The set E can be chosen such that* |*E*| ≤ dim(*C*) *and* |*E*| ≤ | supp(*x*)|+| supp(*Ax*)|.

#### A.3 Polyhedra

Let *P* be a polyhedron, that is, *P* = {*x* ∈ ℝ^*n*^ | *Ax* ≥ *b*} for some *A* ∈ ℝ*m*×*n* and *b* ∈ ℝ*m*. A vector *x* ∈ *P* is *convex-conformally non-decomposable* (ccND) if for all *x*^1^, *x*^2^ ∈ *P* with sign(*x*^1^), sign(*x*^2^) ≤ sign(*x*) and 0 *< λ <* 1, the decomposition *x* = *λx*^1^ + (1 − λ)*x*^2^ implies *x*^1^ = *x*^2^.

Let *R* = {*x* ∈ ℝ^*n*^ | *Ax* ≥ 0} be the recession cone of *P*. A vector *e* ∈ *P* ∪ *R* is

*elementary* (an EV of *P*) if *e* ∈ *P* is ccND or *e* ∈ *R* is cND.

Ultimately, Theorem 43 can be extended to general polyhedral cones. See [19, Theorem 13].

##### Theorem 45.

*Let P* = {*x* | *Ax* ≥ *b*} *be a polyhedron and R* = {*x* | *Ax* ≥ 0} *its recession cone. Every vector x* ∈ *P is a conformal sum of EVs. That is, there exist finite sets E*_0_ ⊆ *R and E*_1_ ⊆ *P of EVs such that*

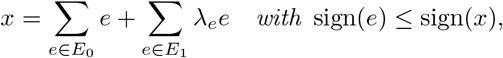

λ_*e*_ ≥ 0, *and* 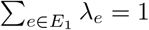. (*Hence*, |*E*_1_| ≥ 1.)

The set E = *E*_0_ ∪ *E*_1_ *can be chosen such that* |*E*| ≤ dim(*P*) + 1 *and* |*E*| ≤ | supp(*x*)| + | supp(*Ax*)| + 1.

### B A minimal derivation of the dynamic growth model

We denote fundamental objects and quantities as follows:

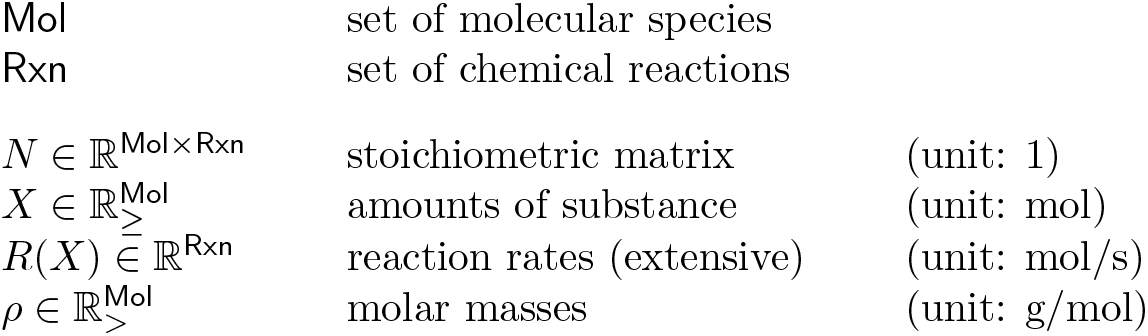

|  |  |  |
| --- | --- | --- |
| Mol | set of molecular species |  |
| Rxn | set of chemical reactions |  |
| $N \in \mathbb{R}^{\text{Mol} \times \text{Rxn}}$ | stoichiometric matrix | (unit: 1) |
| $X \in \mathbb{R}_{\geq}^{\text{Mol}}$ | amounts of substance | (unit: mol) |
| $R(X) \in \mathbb{R}^{\text{Rxn}}$ | reaction rates (extensive) | (unit: mol/s) |
| $\rho \in \mathbb{R}_{>}^{\text{Mol}}$ | molar masses | (unit: g/mol) |

The chemical reactions induce the dynamical system

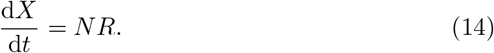

We define dry weight (correctly: dry mass),

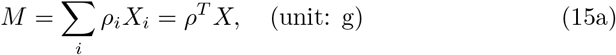

the intensive quantities

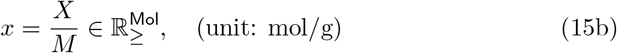

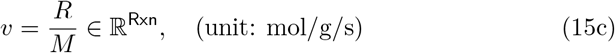

and growth rate

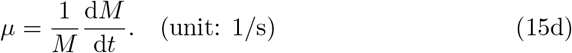

#### Note

We use mass instead of volume to define the “concentrations” *x*, the (intensive) reaction rates *v*, and growth rate *μ*. Thereby, we avoid discussions about “molecular volume”, osmotic pressure, etc. at this point. Moreover, in practice, cellular composition is often given in the unit mol/g (dry weight). Finally, we recall the chain rule (of differentiation),

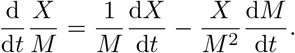

Now, Equations (14), (15), and the chain rule yield the dynamic model of growth.

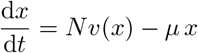

and

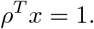

#### Note

The derivation even holds for unbalanced growth.

### C Mathematical issues in [de Groot et al, 2020]

The paper [5] is inspirational from the modeling perspective. Unfortunately, the mathematical treatment is not rigorous. In particular, Definition 1 is flawed, and a crucial intermediate result (regarding the support minimality of EGSs, cf. Proposition 29 in this work) is missing. As a consequence, almost every statement and/or its proof is imprecise or incomplete.

#### Incorrect statements

- Definition 1: 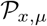 is not a polytope, in general, but a polyhedron.
- Theorem 1: As a consequence of Definition 1, not every element of 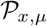 is a convex combination its vertices (EGSs), in general.
- Theorem 3: A condition on the support of *x*_0_ is missing. Obviously, if (*x*_0_)_*i*_ = 0 for some *i*, then there is no neighbourhood of (*x*_0_, *μ*_0_) such that the EGS *α* can be continuously extended. Moreover, the statement has an unnecessary assumption, namely, that the support of the EGS equals its feasible basis.
- Theorem 5: As another consequence of Definition 1, even if the polyhedron 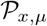 intersects the polyhedron *P*_cons_ determined by more than one inequality constraint, the polytope *P*_EGSs_ generated by the vertices (EGSs) of 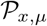 does not intersect *P*_cons_, in general. Hence, feasible solutions (in particular, optimal solutions) are not sums of EGSs, in general.
- Theorem 6: The statement requires the assumption that *P_D_* has full rank. Under the assumption that (*P_D_* − *ϕ*) has full rank, row reduction applied to (*P_D_* − *ϕ*) does not yield 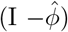 Moreover, certain dimensions do not match. On the one hand, *D* ⊆ 1,… *n* + 1 with |*D*| = *m* + 1 is a feasible basis (of *B* ∈ ℝ^(*m*+1)×(*n*+1)^). On the other hand, *P* ∈ ℝ^*m*×*n*^ is the stoichiometric matrix. Hence, *P_D_* is ill-defined. Finally, the notation 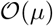 is misleading. In the sum 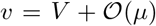, also the “biomass flux mode” *V* is 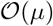.
- Theorems 7 and 8 are based on the incorrect Theorem 6.

#### Incorrect proof

- Theorem 4: Theorem 3 is used without checking the (unnecessary) assumption that the support of the EGS equals its feasible basis. Moreover, the proof is too complicated, in particular, the implicit function theorem is not needed.

